# Anxious and obsessive-compulsive traits are independently associated with valuation of non-instrumental information

**DOI:** 10.1101/768168

**Authors:** Daniel Bennett, Kiran Sutcliffe, Nicholas Poh-Jie Tan, Luke D. Smillie

## Abstract

Aversion to uncertainty about the future has been proposed as a transdiagnostic trait underlying psychiatric diagnoses including obsessive-compulsive disorder and generalised anxiety. This association might explain the frequency of pathological information-seeking behaviours such as compulsive checking and reassurance-seeking in these disorders. Here we tested the behavioural predictions of this model using a non-instrumental information-seeking task that measured preferences for unusable information about future outcomes in different payout domains (gain, loss, and mixed gain/loss). We administered this task, along with a targeted battery of self-report questionnaires, to a general-population sample of 146 adult participants. Using computational cognitive modelling of choices to test competing theories of information valuation, we found evidence for a model in which preferences for costless and costly information about future outcomes were independent, and in which information preference was modulated by both outcome mean and outcome variance. Critically, we also found positive associations between a model parameter controlling preference for costly information and individual differences in latent traits of both anxiety and obsessive-compulsion. These associations were invariant across different payout domains, providing evidence that individuals high in obsessive-compulsive and anxious traits show a generalised increase in willingness-to-pay for unusable information about uncertain future outcomes, even though this behaviour reduces their expected future reward.

---

Uncertainty is an inescapable condition of living in a stochastic and dynamic world, and thus an ever-present feature of the decision problems that humans face each day. Given this ubiquity, one might naively expect that humans would habituate to being uncertain, and that their decisions would therefore be insensitive to the presence of uncertainty. This is not the case; instead, human behaviour is exquisitely sensitive to differences in uncertainty between different courses of action or sources of information. This sensitivity is manifest in economic phenomena such as risk aversion and ambiguity aversion (Allais, 1953; Bernoulli, 1954), in the time course of perceptual decision making on the basis of noisy sensory inputs (Reddi et al., 2003; Smith & Ratcliff, 2004), in selective attention for associative learning (Dayan et al., 2000), and in the weight assigned to unreliable social sources of information (Birnbaum & Mellers, 1983).

Behavioural sensitivity to uncertainty across multiple distinct domains suggests the possibility that humans may hold preferences with respect to uncertainty itself. Some theories postulate that the amount of uncertainty associated with a particular action or cognitive state can itself be appetitive or aversive (Freeston et al., 1994; Krohne, 1993; Sorrentino et al., 1990). Moreover, a trait-level tendency to find uncertainty about future events aversive has been proposed as a transdiagnostic feature of psychiatric illness, especially obsessive-compulsive and anxiety disorders (Carleton, Sharpe, et al., 2007; Carleton et al., 2012; Dugas et al., 2001; McEvoy & Erceg-Hurn, 2016; Miceli & Castelfranchi, 2005; Sarawgi et al., 2013; Tolin et al., 2003). In support of this contention, self-report questionnaire measures of intolerance of uncertainty have been shown to correlate positively with the severity of symptoms of disorders including obsessive-compulsive disorder (OCD), generalised anxiety disorder, and social anxiety (Carleton et al., 2012; Fetzner et al., 2013; Gentes & Ruscio, 2011; McEvoy & Erceg-Hurn, 2016; Sarawgi et al., 2013; Tolin et al., 2003). Aversion to uncertainty might also account for symptoms of these disorders such as excessive reassurance-seeking and compulsive checking (Kobori & Salkovskis, 2013; Lind & Boschen, 2009; Obsessive Compulsive Cognitions Working Group, 1997). In individuals who experience extreme aversion to uncertainty, such behaviours can be rationalised as attempts to gain relief from uncertainty via information-seeking.

These psychiatric theories predict that individuals high in obsessive-compulsive and anxious traits should display increased information-seeking in response to uncertainty. However, behavioural findings in this regard have been extremely mixed. A number of studies have attempted to study information-seeking behaviour in OCD and generalized anxiety disorder (Chamberlain et al., 2007; Fear & Healy, 1997; Foa et al., 2003; Garety et al., 1991; Grassi et al., 2015, 2018; Hauser, Moutoussis, Iannaccone, et al., 2017; Hauser, Moutoussis, NSPN Consortium, et al., 2017; Hezel et al., 2019; Jacobsen et al., 2012; Jacoby et al., 2014; Milner et al., 1971; Pélissier & O’Connor, 2002; Schlier et al., 2016; Toffolo et al., 2013; Volans, 1976; Voon et al., 2017) using either the beads task (Huq et al., 1988; Phillips & Edwards, 1966) or the information sampling task (Clark et al., 2006). Both of these tasks assay preference for *instrumental* information (that is, information which can be used to improve a future decision): in the beads task, participants observe coloured beads drawn sequentially from one of two urns with known colour proportions (e.g., one urn with 85 pink beads and 15 green, and another with 15 pink beads and 85 green). After each bead is drawn, participants choose either to guess which urn is being drawn from or to observe another draw. Similarly, in the information sampling task participants are presented with a 5×5 grid of closed boxes, each of which reveals one of two colours when opened. In this task the goal is to infer which of the two colours is in the majority in the grid, and participants can open as many boxes as they wish prior to making a response. Using these tasks, some studies have found that participants with OCD and those on the compulsivity spectrum request more information prior to making a decision than healthy controls, consistent with greater aversion to uncertainty (Fear & Healy, 1997; Hauser, Moutoussis, Iannaccone, et al., 2017; Hauser, Moutoussis, NSPN Consortium, et al., 2017; Pélissier & O’Connor, 2002; Volans, 1976; Voon et al., 2017). On the other hand, other studies have found either no difference in information-seeking between participants with OCD and controls (Chamberlain et al., 2007; Hezel et al., 2019; Jacobsen et al., 2012), or evidence for *less* information-seeking in OCD than in controls (Grassi et al., 2015, 2018). Results are similarly mixed for anxiety-relevant traits: although there is some evidence that information seeking is increased in social anxiety disorder (Schlier et al., 2016), and correlates with self-reported anxiety (Crockett et al., 2012), other studies have found no alterations in information-seeking in generalised anxiety disorder (Garety et al., 1991; Jacoby et al., 2014).

These inconsistencies appear to challenge the proposal that aversion to uncertainty is an important risk factor for the development of psychopathology (Dugas et al., 1997; Koerner & Dugas, 2006), as well as for cognitive therapies based on these theories that target intolerance of uncertainty in patients with generalized anxiety disorder (Ladouceur et al., 2000; van der Heiden et al., 2012; Wells & King, 2006). However, the mere fact that the information offered to participants in the beads and information-seeking tasks is instrumental introduces two cognitive confounds to these tasks. First, irrespective of aversion to uncertainty, the *economic* value of instrumental information depends not only on one’s uncertainty, but also on the absolute utility or disutility of possible outcomes (Lawrence, 1999; Raiffa & Schlaifer, 1961): the worse a negative outcome, the more valuable is information that may allow one to avoid it. As a consequence, a plausible explanation for stronger information-seeking in those high in anxious and obsessive-compulsive traits is over-estimation of future threat (Obsessive Compulsive Cognitions Working Group, 1997): If potential negative outcomes are seen as more disastrous, as evidence suggests (Paul et al., 2016; Rouel et al., 2018), the instrumental value of information will naturally increase to them as a consequence. Secondly, instrumental information-seeking actions also reduce the *risk* of a future decision. In the beads task, for instance, making a prediction about the urn that is being drawn from is a risky decision problem. By drawing a new bead, the participant effectively chooses to face a new decision problem that is likely to be less risky than the first (assuming the reduction of compound lotteries). A decision to acquire information in this task may therefore be the result of risk aversion in the economic sense of decreasing marginal utility, distinct from intolerance of uncertainty as defined in the psychological and psychiatric literature (Freeston et al., 1994; Krohne, 1993). Given that increased risk and ambiguity aversion have been linked with both anxiety and OCD (Pushkarskaya et al., 2015; Steketee & Frost, 1994; Charpentier et al., 2017), this represents another barrier to interpretation of commonly used information-seeking tasks in terms of aversion to uncertainty.

The present study used a *non-instrumental* information seeking task (Bennett et al., 2016; Brydevall et al., 2018) to compute a behavioural measure of aversion to uncertainty that is not affected by these confounds. In contrast to instrumental information, non-instrumental information reduces one’s uncertainty but cannot be used to alter any future behaviour (for example, information about whether or not a foregone conclusion will occur). Concretely, in our task participants could acquire early information regarding the outcome of a lottery, but this information could not be used to alter the lottery outcome in any way. Willingness to acquire non-instrumental information is a relatively direct measure of aversion to uncertainty, since reducing uncertainty is by definition the only function that non-instrumental information serves. This behaviour is also unconfounded by threat overestimation or risk aversion, since these confounds are induced by the instrumentality of information. Previous research has shown that humans and other animals behave as though non-instrumental information has an intrinsic value, and will even pay an explicit cost (such as a monetary cost, a wait-time cost, or foregone primary reward) to acquire it (Bennett et al., 2016; Blanchard et al., 2015; Bromberg-Martin & Hikosaka, 2009, 2011; Brydevall et al., 2018; Cabrero et al., 2019; Charpentier et al., 2018; Iigaya et al., 2016; Kobayashi et al., 2019; van Lieshout et al., 2018; Vasconcelos et al., 2015), even though this leads to a reduction in future expected reward. In humans, there is marked heterogeneity in the strength of this preference (Bennett et al., 2016), suggesting individual differences in aversion to uncertainty that resemble those proposed in theories of obsessive-compulsion and anxiety (Krohne, 1993; Sorrentino et al., 1990).

In the following study, we investigated the relation between willingness to pay for non-instrumental information in this behavioural task and three subclinical traits related to psychiatric phenomena (obsessive-compulsion, anxiety/negative emotionality, and need for structure/control) as quantified by a targeted questionnaire battery in a general-population sample. A general-population sample allowed us to test a dimensional model of the relation between aversion to uncertainty and psychopathology, in line with dimensional proposals regarding aversion to uncertainty specifically (Carleton et al., 2012), as well as psychopathology more broadly (Haslam et al., 2012; Robbins et al., 2012; Widiger et al., 2019). Our task design also allowed us to extend previous empirical work on valuation of non-instrumental information in two respects. First, we compared willingness to pay for information about lotteries in the gain domain (i.e., a lottery over a monetary gain and a non-gain) versus the loss domain (a lottery over a monetary loss and a non-loss), as well as a mixed gain-loss domain, testing the propositions that the subjective value of non-instrumental information differs between gains and losses and scales with the variance of potential outcomes (Charpentier et al., 2018; Golman & Loewenstein, 2018; Kobayashi et al., 2019). Second, we directly tested a common assumption in the literature, namely that willingness to pay for information and willingness to acquire costless information are related manifestations of a single latent information preference. Since psychiatric symptoms such as reassurance-seeking or checking behaviours are typically associated with significant time or social costs, we revisited this assumption, testing whether subclinical traits were differentially associated with willingness to pay for information versus willingness to acquire costless information.

## Method

### Participants

Participants were 146 members of the general public, recruited via online advertisements and posters displayed at the Parkville campus of the University of Melbourne. All participants were individually tested in controlled laboratory conditions, and participants received a flat rate of AUD $10 compensation, plus their winnings from the behavioural task (mean winnings = $14.32, SD = 1.05). Data was discarded from seven participants who failed an attention check in the behavioural task (detailed below), or who failed to respond on more than 10 percent of choice trials. The final sample consisted of 139 participants (98 female, 40 male, 1 unspecified gender) aged 18 to 35 years (*M* = 23.99, *SD* = 3.77). This sample size marginally exceeded our *a priori* target sample size of 133, which was chosen to provide 80% power to detect a typical effect size in individual-differences research (*r* = .24) (Fraley & Marks, 2007). Participants provided written informed consent, and this study was approved by the Human Research Ethics Committee of the University of Melbourne (ID 164851.6). Research was conducted in accordance with APA ethical standards for human-subject research.

### Materials and Procedure

In a soundproofed behavioural testing booth, each participant completed both a non-instrumental information seeking task and a battery of computerised questionnaires. The order of completion of the questionnaire battery and behavioural task was counterbalanced across participants

### Behavioural task

Participants completed nine blocks of a computerised non-instrumental information-seeking task, each consisting of 20 trials. Each trial of the task comprised a lottery in which one of two equiprobable monetary outcomes occurred after a six-second delay. During this delay period, participants could choose either to observe an informative stimulus that perfectly predicted the outcome of the lottery, or to observe a non-informative stimulus that was perceptually identical to the informative stimulus but unpredictive of the lottery outcome (Figure 1). Both stimuli were arrays of five cards presented face-down; when revealed, cards could be either red or black. The chosen stimulus was then revealed at a rate of one card per second. In the informative stimulus, a majority of black cards predicted a win, and a majority of red cards a loss. In the non-informative stimulus, the majority card colour was unrelated to the lottery outcome.

**Figure 1.**
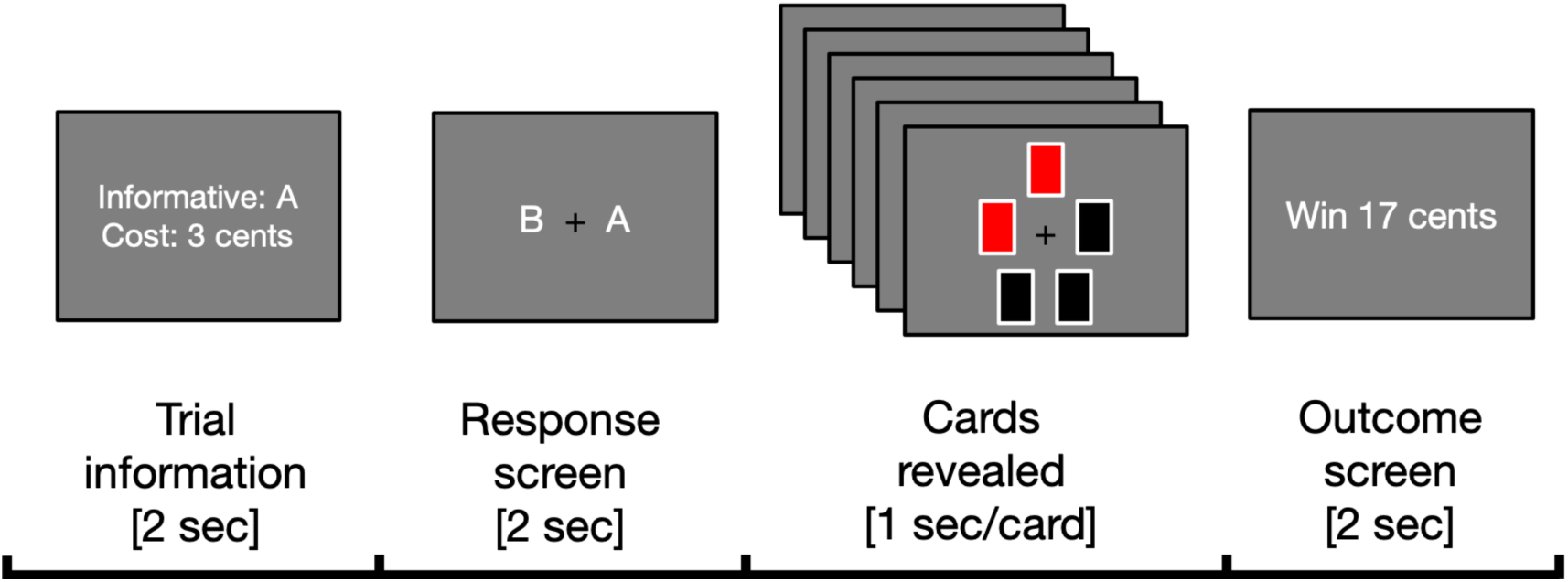
Trial schematic. Participants were presented with a choice between stimulus A and stimulus B, where either A or B could be informative on any given trial. Before making a choice they were informed which stimulus was informative, and the cost associated with this informative stimulus.

Participants completed three blocks in each of three outcome domains: a gain-domain condition (win = +20c, loss = 0c), a loss-domain condition (win = 0c, loss = −20c), and a high-variance condition (win = +20c, loss = −20c). In each trial, one of three costs was applied to choices to view the informative stimulus (0, 1, or 3c). Different payout domains (i.e., gain domain, loss domain, mixed domain) were presented in different blocks, and block order was randomised subject to the constraint that the same payout domain could not be presented in consecutive blocks. Each trial was randomly associated with either a 0, 1, or 3-cent cost on the informative stimulus (20 trials per cost per domain; within a domain, trials in each cost condition were pseudo-randomly distributed across the three blocks assessing information preference in that domain). This cost was always paid if participants chose to observe the informative stimulus, regardless of whether the lottery outcome was a win or a loss. Because the lottery was independent of participants’ choice between the informative and the non-informative stimulus, paying a cost to observe the informative stimulus is always sub-optimal (since it equates to choosing a lottery with a lower expected value). Participants began the experiment with an AUD $15 endowment, which could be used to pay for information and to which lottery outcomes were added or subtracted.

Participants completed five practice trials and proceeded to testing when experimenters were confident that they understood task instructions. The task was presented using the Psychophysics Toolbox (Brainard & Vision, 1997) on a Macintosh Mini desktop computer running Matlab (release 2012b). Participants entered responses using the left and right arrow keys and, as in previous studies using this task, an attention check was used to ensure that participants maintained attention to the stimulus after they had made a choice. Specifically, on ten percent of trials (two per block), one of the cards in the chosen stimulus was revealed to contain a white cross rather than the expected red or black card. When this happened, participants were instructed to respond by pressing any key on the keyboard within 1.5 seconds. If they failed to do so, one dollar was deducted from their endowment. Applying the same exclusion criterion as in previous studies (Bennett et al., 2016; Brydevall et al., 2018), we excluded participants’ data from further analysis if they failed to respond to more than two catch trials in total across the task.

### Questionnaire battery

Participants completed a targeted questionnaire battery consisting of nine distinct self-report scales (see Method for details), designed to quantify three trait-level domains of individual difference that are theoretically linked to aversion to uncertainty: obsessive-compulsion, anxiety/negative emotionality, and need for structure/control (Berenbaum et al., 2008; Koerner et al., 2017; Shihata et al., 2016; Webster & Kruglanski, 1994). These three constructs were chosen *a priori*, and we included multiple scales measuring each to ensure that our interpretation of results was not restricted to any one measure of each construct. Consequently, we did not analyse individual scales directly. Instead, we performed dimensionality reduction separately for each construct using a principal component analysis (see below) to extract a per-participant measure that was entirely based upon the shared variance among distinct measures of the construct, thereby reducing measurement error.

In addition, separately from this factor-analytic method, participants also completed the 12-item Intolerance of Uncertainty (IU-12) scale (Carleton, Norton, et al., 2007). This scale is the most commonly used self-report measure of intolerance of uncertainty in the literature, and allowed us to assess the relations between performance on the non-instrumental information-seeking task and self-report of intolerance to uncertainty. We therefore did not include the IU-12 scale in the dimensionality reduction analysis but analysed it separately. The questionnaire battery completed by participants also included several measures of other Big Five personality constructs for a separate student research project. These additional personality measures were unrelated to the research questions of the present study and are therefore not reported here.

The entire self-report battery comprised 86 items. Unless otherwise specified below, for all scales participants rated the accuracy with which a descriptive statement described them (e.g., “I wash my hands more than is necessary”) using a 5-point Likert scale (anchors: “Very Inaccurate”, “Very Accurate”). The order of presentation of questionnaires within the battery was pseudo-randomised across participants. The questionnaire instruments included in the battery were as detailed below, organised by the latent trait that each instrument was designed to quantify.

#### Obsessive-compulsive traits

Scales chosen to assess obsessive-compulsive traits were:

1. the 12-item International Personality Item Pool (IPIP; Goldberg et al., 2006) version of the Obsessive-Compulsive Inventory (Revised) (Foa et al., 2002) (sample item: “I get upset if objects are not arranged properly”).
2. the 10-item rigid perfectionism scale (designed to measure personality traits related to OCD at subclinical levels) from the Personality Inventory for DSM-5 (Krueger et al., 2012). Participants rated the extent to which a descriptive statement was true of them (e.g., “I check things several times to make sure they are perfect”) using a 4-point Likert scale (anchors: (anchors: “Very False or Often False” and “Very True or Often True”).
3. the 10-item IPIP version of the Need for Order and Cleanliness scale (Foa et al., 1998) (sample item: “I want everything to be ‘just right’”).

#### Need for organisation and control

Scales chosen to assess need for structure and control were:

(4) the 4-item organisation subscale of the Big Five Inventory-2 (BFI-2; (Soto & John, 2017). Participants were asked to rate their agreement with descriptive statements (e.g., “I am a person who is systematic, likes to keep things in order”) on a 5-point Likert scale (anchors: “Disagree strongly”, “Agree strongly”).
(5) the 10-item IPIP version of the orderliness subscale of the Big Five Aspects Scale (BFAS; DeYoung, Quilty, & Peterson, 2007) (sample item: “I want every detail taken care of”).

#### Anxiety and negative emotionality

Scales chosen to assess anxiety and negative emotionality were drawn from the BFI-2 and the BFAS:

(6) the 4-item anxiety subscale of the BFI-2. Participants were asked to rate their agreement with descriptive statements (e.g., “I am someone who worries a lot”) on a 5-point Likert scale (anchors: “Disagree strongly”, “Agree strongly”.)
(7) the 4-item emotional volatility subscale of the BFI-2. Participants were asked to rate their agreement with descriptive statements (e.g., “I am a person who is temperamental, gets emotional easily”) on a 5-point Likert scale (anchors: “Disagree strongly”, “Agree strongly”.)
(8) the 10-item IPIP version of the withdrawal subscale of the BFAS (sample item: “I am filled with doubts about things).
(9) the 10-item IPIP version of the volatility subscale of the BFAS (sample item: “I get upset easily”).

### Data Analysis

All frequentist analyses controlled for Type 1 errors at α = .05, and all reported *p*-values are two-tailed. Mixed-effects logistic regression analyses were conducted using the lme4 package in R (Bates et al., 2015), with random intercepts for participants and fixed effects selected according to a maximal-to-minimal-that-converges procedure (Meteyard & Davies, 2020). The *p*-values for coefficient estimates in mixed-effects models were estimated using asymptotic Wald tests.

### Principal component analysis

To perform dimensionality reduction on the questionnaire data, we conducted a separate principal component analysis on scale totals within each of the three *a priori* trait domains (obsessive-compulsion, need for structure/control, and anxiety/negative emotionality). Factor scores for the first principal component in each of these domains was used to quantify individual differences on this trait.

### Computational modelling of choice data

We formulated and compared 13 competing computational models of participants’ choice behaviour using a hierarchical Bayesian approach. Different models tested distinct hypotheses about how participants calculated the value of the informative stimulus; specifically, models differed in terms of their assumptions about two factors: (a) the source of the information’s value, and (b) the functional form of the price elasticity of information.

For the first factor, models assumed either that information preference covaried with the mean lottery outcome (models 3, 7, 11), with the variance of lottery outcomes (models 4, 8, 12), both (models 5, 9, 13), or neither (models 1, 2, 6, 10). Formal comparison of these model families allowed us to test two hypotheses regarding information preference about uncertain future outcomes. First, models incorporating a modulation of information preference mean lottery outcomes test the hypothesis that information about future outcomes in the gain domain is preferred more than information about outcomes in the loss domain (Charpentier et al., 2018; Golman et al., 2017). Second, models incorporating a modulation of information preference by the variance of lottery outcomes test the hypothesis that this variance is a marker of the importance of information, and therefore a determinant of preference for information (Golman & Loewenstein, 2018).

Differences between models on the second factor, price elasticity, were introduced to capture our empirical finding that preference for information was not positively correlated between the zero-cost and the non-zero-cost conditions (see Results). Since price elasticity— defined as the rate at which preference for some good changes as its price changes—is a mechanism which can account for this lack of a positive correlation, we tested three families of models to account for this effect. In the first family, the price elasticity of information was constant (models 2-5); in the second, price elasticity varied linearly across participants (model 6-9); in the third, price elasticity of information was variable and non-linear (models 10-13). Importantly, although we did not have *a priori* hypotheses for which of these models would best explain the data, we independently verified that the lack of correlation between costly and cost-free information was also present in previously collected datasets (see Results). This gives us confidence that we are not over-fitting models to noise in choice data in the present study.

All models assumed that participants’ choices were distributed according to a softmax function, such that the probability of choosing the informative stimulus (denoted *I*) was proportional to the difference in subjective value between it and the non-informative stimulus (denoted *N*). This amounts to a sigmoidal choice rule, where the steepness of the sigmoid was controlled by the inverse temperature parameter *β* such that higher values of *β* corresponded to a steeper sigmoid and therefore more deterministic choices:

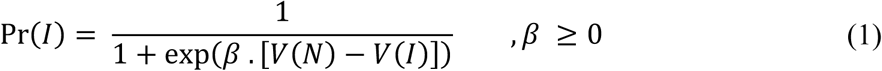

Note that *β* was fixed to a value of 1 for models 6-10, for reasons discussed below.

Across all models, the value of the non-informative stimulus *V*(*N*) was simply defined as the expected value of the trial lottery *L:*

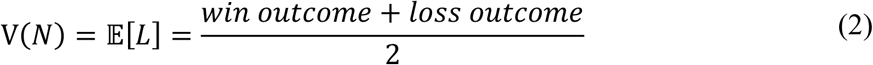

As a result, differences between models resulted from differences in how they assumed the value of the informative stimulus *V*(*I*) was computed. As described above, with the exception of the null model (model 1), models differed on two dimensions:

A. how different features of the lottery outcome were assumed to affect the value of information, and
B. how preference for information were assumed to vary with changes in the cost of information (i.e., the price elasticity of information).

For the best-fitting model, point estimates of each parameter were calculated per participant as the median of the participant-level posterior distribution.

#### Model 1: Null model

The simplest null model (model 1) embodied the null hypothesis that participants did not assign any intrinsic value to information. In this case, the value of the informative stimulus is simply equal to the expected value of lottery minus the cost *c* of observing the informative stimulus:

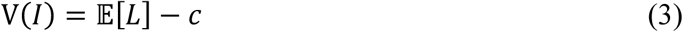

#### Model 2: Baseline information preference, constant price elasticity

Model 2 assumed that the value of information was equal to a constant (the parameter *ϕ*) across all lottery domains:

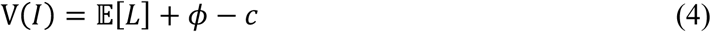

Since information’s value in this model is a constant, model 2 is incapable of predicting any differences in information preference between the gain, loss, and mixed domains.

#### Model 3: Mean-dependent information preference, constant price elasticity

Model 3 assumed that the value of information was equal to a constant plus a mean-dependent component that varied according to the expected value of the lottery (modulated by the parameter *k*_*mean*)_:

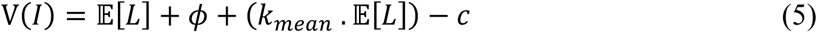

Unlike model 2, model 3 can predict differences in information preference between the gain, loss, and mixed domains. Specifically, for *K*_*mean*_> 0, the participant prefers information most in the gain domain and least in the loss domain; for *K*_*mean*_< 0, the participant prefers information most in the loss domain. When *K*_*mean*_= 0, model 3 reduces to model 2.

#### Model 4: Variance-dependent information preference, constant price elasticity

Model 4 assumed that the value of information was equal to a constant plus a variance-dependent component that varied according to the variance of the lottery (modulated by the parameter *k*_*var*_):

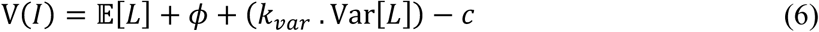

Model 4 predicts differences between the mixed domain and both of the gain and loss domains. Specifically, for *K*_*var*_ > 0, the participant prefers information more strongly in the mixed domain than in either the gain or the loss domain, and vice versa for *K*_*var*_ < 0. Unlike model 3, this model is incapable of predicting differences in information-seeking between the gain domain and the loss domain. When *K*_*var*_ = 0, model 4 reduces to model 2.

#### Model 5: Mean- and variance-dependent information preference, constant price elasticity

Model 5 combines model 3 and 4 by assuming that the value of information was equal to a constant plus both a mean-dependent component and a variance-dependent component:

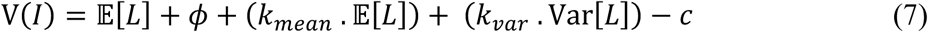

#### Model 6: Baseline information preference, linear price elasticity

Similar to model 2, model 6 assumed that the value of information was equal to a constant across all lottery domains. Unlike model 2, model 6 allowed for individual differences in the steepness with which the value of information fell off as its cost increased:

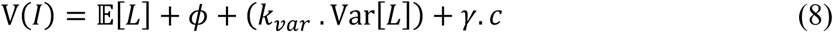

In Equation 8, the parameter *γ* controls the price elasticity of information preference. *γ* is typically negative, such that the subjective value of the informative stimulus decreases as its cost increases (as is intuitive). When *γ* is close to zero, participants’ preference for information is relatively *inelastic*, in the sense that changes in the cost of the informative stimulus have a relatively small effect on participants’ willingness to acquire information. As *γ* becomes more negative, preference for information becomes more elastic, such that information-seeking declines rapidly when a cost is placed on the informative stimulus. Under this framework, model 2 is a special case of model 6 in which *γ* is fixed at −1.

We note that allowing for individual differences in the price elasticity of information introduced an issue of parameter identifiability between the softmax inverse temperature parameter *β* and the price elasticity parameter *γ*. Intuitively, this is because the two parameters can have equivalent effects on choice stochasticity; for example, either a large negative *γ* or a large positive *β* will produce a sigmoidal choice rule that approaches a step function. For this reason (and to avoid divergent transitions resulting from parameter unidentifiability during model fitting), *β* was fixed to a value of 1 in model 6-10.

#### Model 7: Mean-dependent information preference, linear price elasticity

Like model 3, model 7 assumed that the value of information was equal to a constant plus a mean-dependent component that varied according to the expected value of the lottery. Like model 6, model 7 allowed for individual differences in the (linear) price elasticity of information preference:

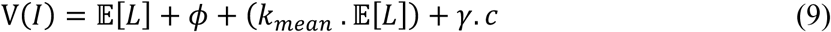

#### Model 8: Variance-dependent information preference, linear price elasticity

Like model 4, model 8 assumed that the value of information was equal to a constant plus a variance-dependent component that varied according to the variance of the lottery. Like model 6 and 7, model 8 allowed for individual differences in the (linear) price elasticity of information preference:

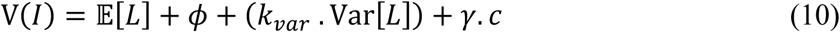

#### Model 9: Mean- and variance-dependent information preference, linear price elasticity

Like model 5, model 9 assumed that the value of information was equal to a constant plus both a mean-dependent component and a variance-dependent component. Like Models 6-8, model 9 allowed for individual differences in the (linear) price elasticity of information preference:

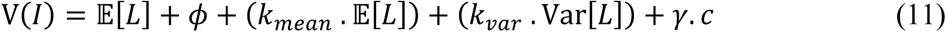

#### Model 10: Baseline information preference, non-linear price elasticity

Like models 2 and 6, model 10 assumed that the value of information was equal to a constant across all lottery domains. However, model 10 assumed the price elasticity of information varied non-linearly as a function of the cost placed on information (thereby giving this model the capacity to explain the finding that preference for information in the zero-cost condition was uncorrelated with preference for information in the non-zero-cost conditions). To avoid making untestable assumptions about the functional form of price elasticity (given that we only had three price points to estimate the shape of the elasticity function), this non-linearity was implemented in the model by allowing the baseline information preference parameter *ϕ* to take on a different value in the zero-cost condition and in the non-zero-cost conditions:

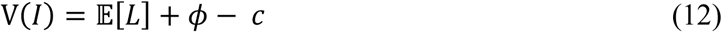

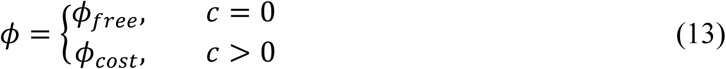

Both *ϕ*_*free*_ and *ϕ*_*cost*_ were free parameters that were allowed to vary across participants. Formally, this is equivalent to fitting a piecewise linear price elasticity function; unlike the linear price elasticity function used in models 6-9, this piecewise linear formulation does not introduce a trade-off with the softmax inverse temperature parameter, and *β* was therefore permitted to vary freely across participants in model 10 (as well as models 11-13).

#### Model 11: Mean-dependent information preference, non-linear price elasticity

Like model 3, model 11 assumed that the value of information was equal to a constant plus a mean-dependent component that varied according to the expected value of the lottery (Equation 5). Like model 10, this model also assumed that the price elasticity of information varied as a non-linear function of the cost placed on information (Equation 13).

#### Model 12: Variance-dependent information preference, non-linear price elasticity

Like model 4, model 12 assumed that the value of information was equal to a constant plus a variance-dependent component that varied according to the variance of the lottery (Equation 6). Like models 10 and 11, this model also assumed that the price elasticity of information varied as a non-linear function of the cost placed on information (Equation 13).

#### Model 13: Mean- and variance-dependent information preference, non-linear price elasticity

Like models 5, model 10 assumed that the value of information was equal to a constant plus both a mean-dependent and a variance-dependent component (Equation 7). Like models 10-12, this model also assumed that the price elasticity of information varied as a non-linear function of the cost placed on information (Equation 13).

### Model fitting

We fit models to data within a hierarchical Bayesian framework, using Hamiltonian Monte Carlo as implemented in Stan (Carpenter et al., 2017) to sample from the joint posterior of all parameters. Four separate chains with randomised start values each took 1500 samples from the posterior. The first 1000 samples from each chain were discarded to prevent dependence on start values, resulting in 2000 post-warmup samples from the joint posterior being retained. 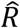 for all parameters was less than 1.1, indicating good convergence between chains, and there were no divergent transitions in any chain. All participant-level parameters were assumed to be drawn from group-level Gaussian hyperprior distributions whose means and standard deviations were estimated freely from the data. In accordance with standard practice in Stan for sampling from complex hierarchical models, parameters were sampled using non-centred parameterisations (all parameters were sampled separately from a unit normal before being transformed to the appropriate range). To prevent negative values the *β* parameter was transformed using the cumulative normal distribution to lie in the range 0-20.

Model comparison was performed using the Watanabe-Akaike Information Criterion (WAIC; Watanabe, 2010), a statistic for comparing models fit with hierarchical Bayesian methods. Like other information criteria (e.g., AIC, BIC, DIC), it selects models according to their goodness-of-fit to data (marginal likelihood, estimated as the mean log-likelihood of data across posterior samples), minus a penalty for the model’s effective complexity (estimated as the variance of the log-likelihood across posterior samples), such that more parsimonious models are favoured over more complex ones (Gelman et al., 2014). We calculated the difference in WAIC between all models, and regarded the best-fitting model as credibly better than its competitors if the standard error of this difference did not overlap with zero.

## Results

### Behavioural Results

Figure 2A presents choice behaviour across payout domains and costs. In line with previous findings (Bennett et al., 2016; Brydevall et al., 2018), on average participants displayed an above-chance preference for observing the informative stimulus when it was free (*p* < .001, Wilcoxon signed-rank test), and displayed a strong modulation of choice behaviour by the cost of information (*β* = −0.90, *p* < .001, mixed-effects logistic regression). Participants also displayed a weaker overall preference for information in the loss domain relative to both the gain domain (*β* = −0.32, *p* < .001) and the mixed domain (*β* = −0.25 *p* < .001). In addition, there was a significant negative effect of trial number (*β* = −0.01, *p* < .01), indicating that participants tended to choose the informative stimulus less as trial number increased within each block. This might be the result of epistemic satiation, such that preference for information about a lottery declines as more and more information is acquired over the course of a block. No other effects were statistically significant (all *p* > .2).

**Figure 2.**
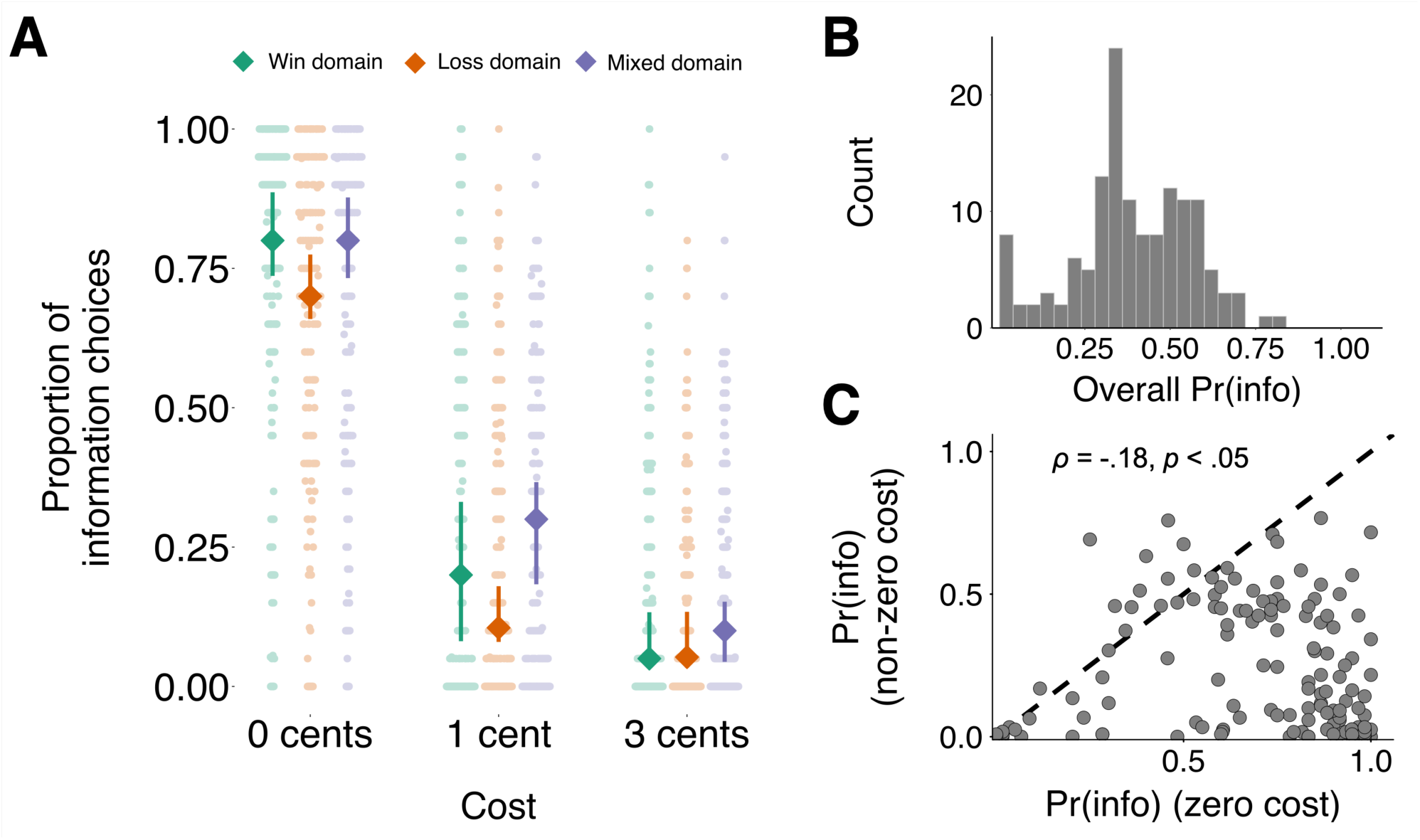
(A) Mean informative stimulus choice proportion as a function of domain and cost. Diamond markers denote condition medians, and error bars denote the bootstrapped 95% confidence interval of the median. Background circular markers denote condition means for individual participants. (B) Participant histogram of informative stimulus choice proportions (denoted Pr(info)) marginalising across cost and lottery domain. (C) Informative stimulus choice proportion for zero-cost trials (horizontal axis) versus non-zero cost trials (vertical axis). The large majority of participants’ choices fall beneath the diagonal (dashed line) in this plot, indicating that they chose information more frequently when it was free than when it was costly. The negative association between these two variables indicates that participants who chose information more frequently when it was free chose information less frequently when it was costly.

Importantly, although information preference was weaker in the loss domain relative to the gain domain, participants did not avoid information in the loss domain. Indeed, when information was free in the loss domain, participants still displayed a clear above-chance preference for observing the informative stimulus (*p* < .001, Wilcoxon signed-rank test).

Across participants, the most common choice pattern was to choose the informative stimulus when it was available at no cost, but to prefer the non-informative stimulus when information was costly. However, as in previous research using this task (Bennett et al., 2016; Brydevall et al., 2018), we observed strong individual differences in patterns of information preference (Figure 2B). These individual differences were consistent across different payout conditions (see Supplementary Table 1), suggesting that the latent cognitive processes underlying information valuation were consistent across gain, loss, and mixed domains.

When examining choices across different cost conditions, however, we found that preference for information was *negatively* correlated between no-cost information trials and trials in which a monetary cost was placed on information (Spearman *ρ* = −.18, *p* < .05; see Figure 2C). In other words, participants who chose information more frequently when it was free chose information *less* when it came at a cost. This negative correlation (which has not previously been documented in the literature) is striking because we had expected to observe a positive correlation, in line with a shared underlying process of information valuation across cost conditions. Indeed, there was a significant positive correlation between preference for information between the 1-cent cost condition and in the 3-cent condition (Spearman *ρ =* 0.85, *p* < .001), suggesting that the negative correlation between costly and costless information preference was not simply an artefact of response noise during choice. Moreover, reanalysis of data from two previously published experiments using a prior variant of the non-instrumental information seeking task (Bennett et al., 2016; Brydevall et al., 2018) showed the same negative correlation (see Supplementary Figure 1). Taken together, results suggest that participants adopted a qualitatively different behavioural strategy in trials where information was available at a cost compared to trials where it was available for free.

As discussed in the Methods, it is possible to account for this apparent strategy difference if we assume that, as well as differing in their overall preference for information, participants also differed in the *price elasticity* of their preference for information. In microeconomics, price elasticity quantifies the degree to which demand for some good changes as the price of that good increases (Mankiw, 1998). Demand for a good is said to be highly *elastic* if it changes rapidly with changes in price, or *inelastic* if it changes little with changes in price. In the non-instrumental information seeking task, choosing information frequently when it was free, but not when it came at a price (the most common pattern of choices across participants) would correspond positive information preference but high price elasticity of information. By contrast, a participant with the same information preference but low price elasticity of information would choose information at a high rate across all three cost conditions. Below, we tested this price-elasticity hypothesis by implementing it in a formal computational model of behaviour (see Computational Modelling Results).

Finally, we investigated whether participants’ preferences for information was modulated by trial-by-trial outcomes associated with the informative stimulus. We reasoned that this pattern would be likely to manifest as a win-stay lose-shift pattern, such that *win* outcomes after viewing the informative stimulus made participants more likely to view the informative stimulus on the subsequent trial, and vice versa for a loss outcome after viewing the informative stimulus. Although this analysis revealed a trend in this direction (*β* = 0.08, *p* = .09; mixed-effect logistic regression), this effect was not statistically significant at α = .05.

### Questionnaire Results

Correlations between the individual scales included in the questionnaire battery are presented in Supplementary Table 2.

Table 1 presents the correlation matrix between the three latent trait constructs and the IU-12 scale (see Supplementary Figure 3 for scatterplots corresponding to each of these correlations). With the exception of the anxiety/negative emotionality factor and the structure/control factor, all trait measures were significantly positively correlated with one another. In particular, all three of the latent factors derived from our questionnaire battery were positively correlated with self-reported intolerance of uncertainty. This gives us confidence that our factor-analytic quantification of these three factors was valid since, as described above, each of these three domains have previously been theoretically linked to aversion to uncertainty in the context of psychopathology.

**Table 1:**
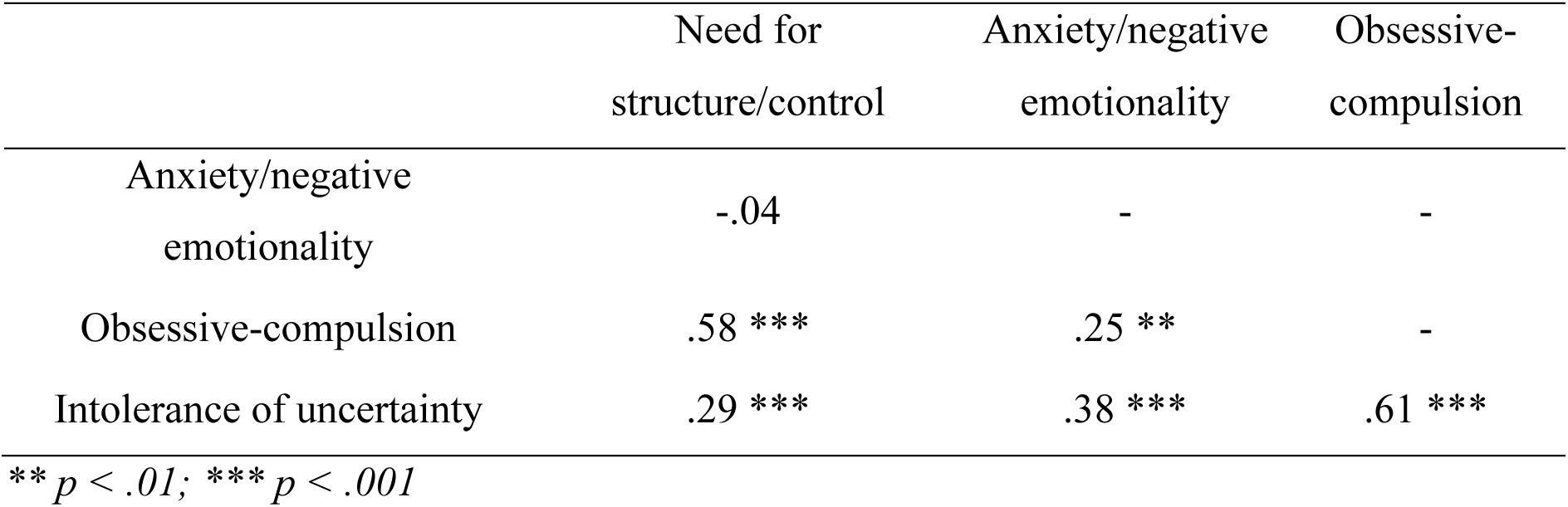
Pearson correlation matrix for factors and scales in self-report battery (N = 139).

|  | Need for<br>structure/control | Anxiety/negative<br>emotionality | Obsessive-compulsion |
| --- | --- | --- | --- |
| Anxiety/negative<br>emotionality | -.04 | - | - |
| Obsessive-compulsion | .58 *** | .25 ** | - |
| Intolerance of uncertainty | .29 *** | .38 *** | .61 *** |

\*\* $p < .01$ ; \*\*\* $p < .001$

### Computational Modelling Results

Of the models that we considered, model 13 (information preference modulated by both lottery mean and variance, non-linear price elasticity of information) provided the best overall fit. Full model comparison results are presented in Table 2.

**Table 2:** Model comparison statistics. ΔWAIC denotes the difference between the WAIC of each model and the WAIC of the best-fitting model.

| Model number | Type of information preference | Type of price elasticity | $n$ parameters per participant | WAIC | $\Delta WAIC$ (S.E.M.) |
| --- | --- | --- | --- | --- | --- |
| 1 | Null | Null | 1 | 25021.58 | 5580.87 (126.64) |
| 2 | Constant | Constant | 2 | 21079.44 | 1638.73 (100.30) |
| 3 | Mean-dependent | Constant | 3 | 20862.93 | 1422.21 (96.43) |
| 4 | Variance-dependent | Constant | 3 | 21037.90 | 1597.19 (99.70) |
| 5 | Mean-dependent and variance-dependent | Constant | 4 | 20822.51 | 1381.80 (95.89) |
| 6 | Constant | Linear | 2 | 20965.83 | 1525.11 (99.00) |
| 7 | Mean-dependent | Linear | 3 | 20660.92 | 1220.21 (94.87) |
| 8 | Variance-dependent | Linear | 3 | 20913.35 | 1472.63 (97.69) |
| 9 | Mean-dependent and variance-dependent | Linear | 4 | 20614.12 | 1173.41 (93.63) |
| 10 | Constant | Non-linear | 3 | 19850.52 | 409.80 (40.03) |
| 11 | Mean-dependent | Non-linear | 4 | 19516.33 | 75.62 (19.21) |
| 12 | Variance-dependent | Non-linear | 4 | 19766.62 | 325.90 (34.95) |
| 13 | Mean-dependent and variance-dependent | Non-linear | 5 | 19440.72 | - |

NB: Both WAIC and $\Delta WAIC$ are presented on a deviance scale, such that lower values indicate better relative model fit.

Furthermore, since the statistics presented in Table 2 only give a measure of the relative (rather than absolute) goodness of fit of models, we also examined a posterior predictive check of the best-fitting model. This posterior predictive check (Figure 3) demonstrates a good correspondence between participants’ observed choices and the model’s predictions (for comparison, posterior predictive checks for the other 12 models are presented in Supplementary Figure 2).

**Figure 3.**
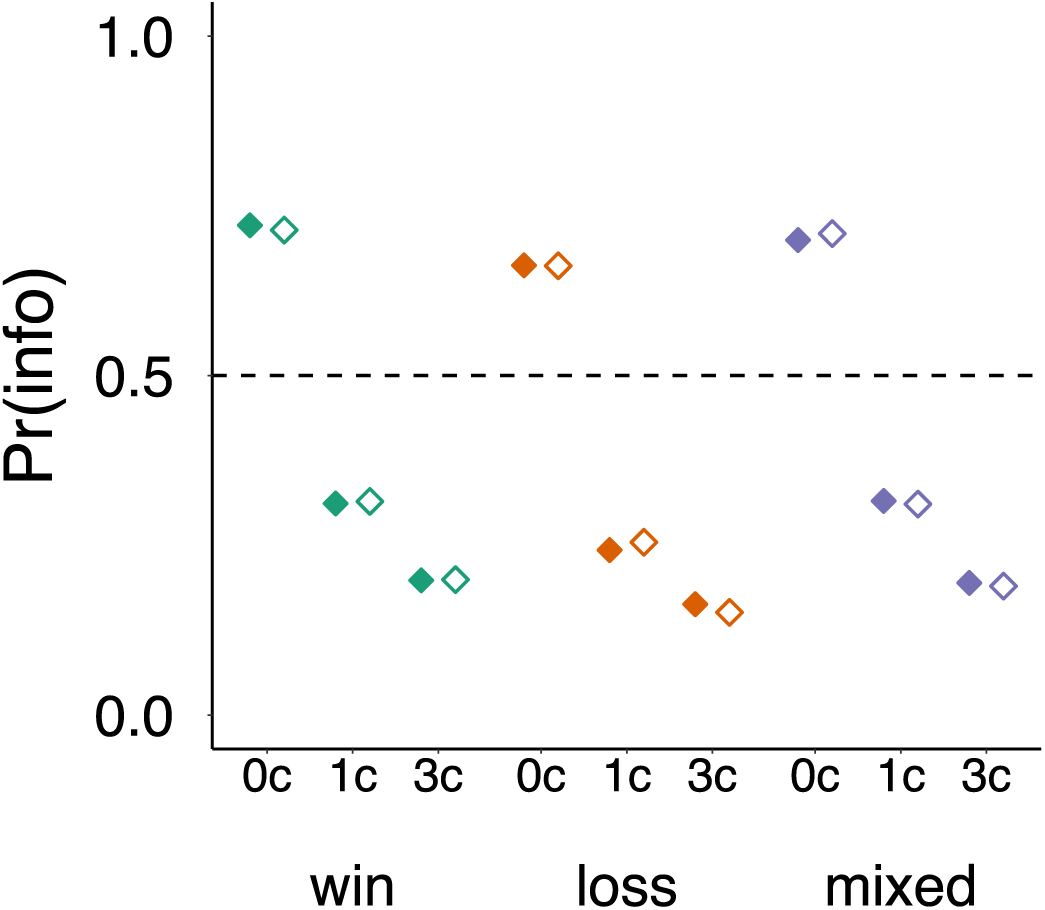
Posterior predictive check for the best-fitting model (model 13). Data are observed (filled markers) and predicted (unfilled markers) mean informative-stimulus choice proportions across payout domains and costs. There was a close correspondence between choice proportions predicted by the model and those observed in the data, indicating good absolute model fit. Pr(info) = proportion of informative stimulus choices.

Model 13 assumed that the price elasticity of information varied non-linearly as a function of cost. This was implemented in the model by assuming that participants had a different underlying information preference for costless versus costly information (respectively quantified by separate baseline information preference parameters, *ϕ*_*free*_ and *ϕ*_*cost*_). For *ϕ*_*free*_ (Figure 4A), the mean of the group-level distribution had a posterior median of 2.01 (95% highest density interval (HDI) [1.39, 2.68]), with a median standard deviation of 3.54 (95% HDI [2.95, 4.23]). Since positive values of this parameter correspond to appetitive valuation of information, this result is in line with the clear preference exhibited by participants for information in the zero-cost condition. For *ϕ*_*cost*_ (Figure 4B) the mean of the group-level distribution had a posterior median of −1.73 (95% HDI [-2.51, −1.06]), and the standard deviation of the group-level distribution had a posterior median of 3.72 (95% HDI [3.10, 4.53]), indicating a preference against costly information on average, but strong individual differences across participants.

**Figure 4.**
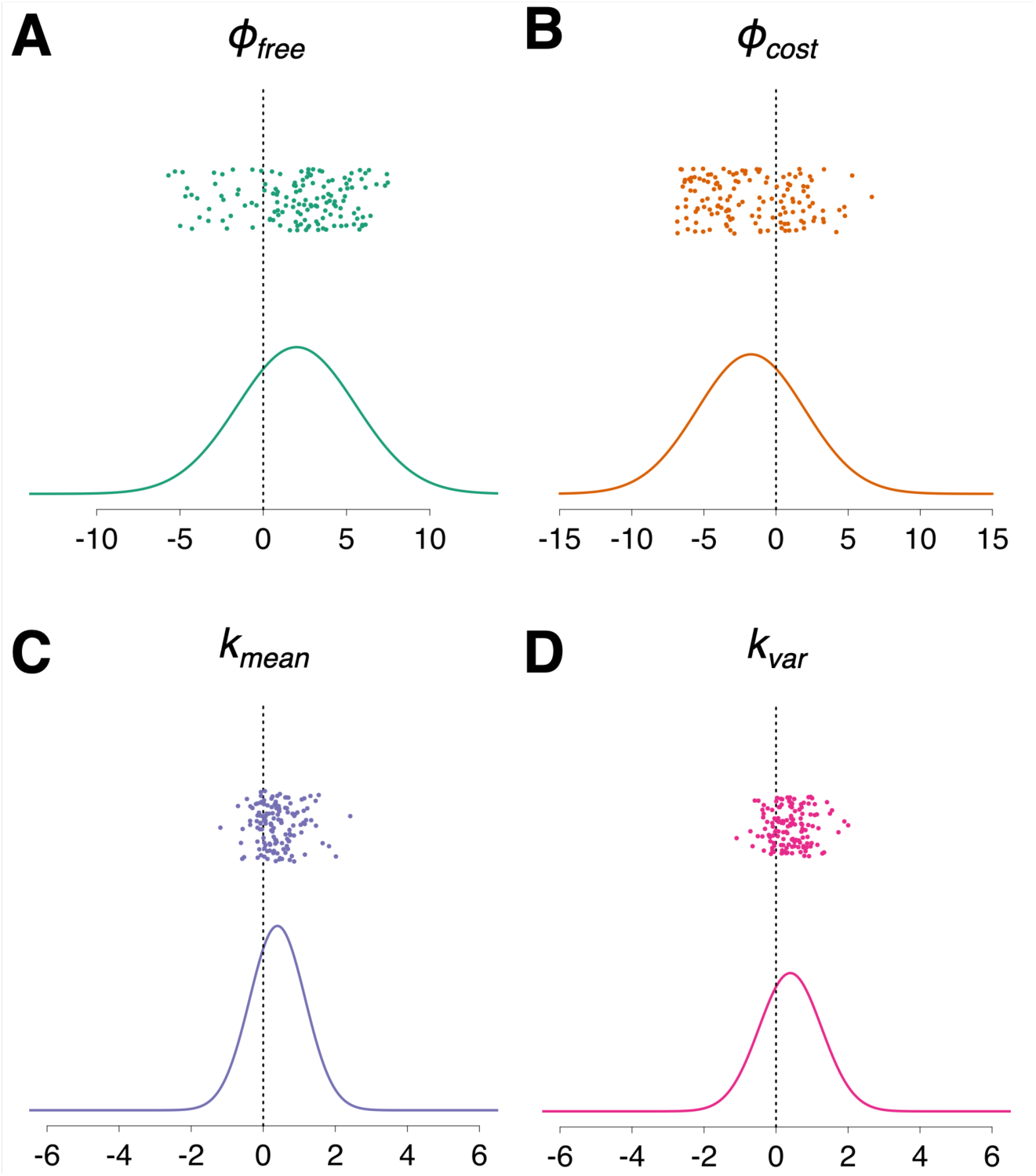
Individual parameter estimates and group-level parameter distributions for the best-fitting computational model. For each parameter, this plot visualises the Gaussian distribution that is implied by the medians of the posterior distributions of the group-level mean and standard deviation. Parameter estimates for individual participants (posterior medians) are presented above each distribution. In green (A): *ϕ*_*free*_ (baseline preference for information in zero-cost trials). In orange (B): *ϕ*_*cost*_ (baseline preference for information in non-zero-cost trials. In purple (C): *k*_*mean*_ (modulation of information preference by lottery outcome mean). In pink (D): *k*_*var*_ (modulation of information preference by lottery outcome variance). In each plot the vertical dashed line represents zero.

Model 13 also allowed for information valuation to vary according to the mean and the variance of the lottery outcome (quantified by the parameters *k*_*mean*_ and *k*_*var*_ respectively). For *k*_*mean*_, the mean of the group-level distribution had a posterior median of 0.39 (95% HDI [0.22, 0.57]). A positive group-level median value of *k*_*mean*_ indicates that, on average, participants sought information more as the expected value of the trial lottery increased. For *k*_*var*_, the mean of the group-level distribution had a posterior median of 0.40 (95% HDI [0.16, 0.63]), in line with a preference for information that increased with lottery variance. The mean of the group-level distribution of the inverse temperature parameter *β* was 0.50 (95% HDI [0.42, 0.59]).

Within-participant parameter correlations are presented in Table 3 (see Supplementary Figure 4 for scatterplots corresponding to each correlation). In line with patterns observed in choice data, there was no correlation between *ϕ*_*free*_ (strength of preference for information in zero-cost trials) and *ϕ*_*cost*_ (strength of preference for information in non-zero-cost trials).

**Table 3:** Pearson correlation matrix of model parameter estimates across participants. N = 139.

| | $\phi_{free}$ | $\phi_{cost}$ | $k_{mean}$ | $k_{var}$ |
| --- | --- | --- | --- | --- |
| $\phi_{cost}$ | -.03 | - | - | - |
| $k_{mean}$ | .05 | .28 *** | - | - |
| $k_{var}$ | .12 | .04 | .09 | - |
| $\log(\beta)$ | .12 | -.38 *** | .02 | .02 |

\*\*\* $p < .001$

There was, however, a significant positive correlation between *ϕ*_*cost*_ and *k*_*mean*_. *k*_*mean*_ quantifies the extent to which participants’ preference for information was modulated by the mean of the lottery outcomes (positive *k*_*mean*_: gain domain preference > mixed domain preference > loss domain preference; vice versa for negative *k*_*mean*_). This positive correlation therefore indicates that participants who paid more for information in costly trials were also more likely to prefer information in the gain domain to information in the loss domain.

Lastly, a negative correlation between *ϕ*_*cost*_ and the logarithm of *β* (the inverse temperature of the softmax function) suggests that participants who paid more for information in costly trials may have also made less deterministic choices in general. In general, however, correlations between other model parameters and *β* should be interpreted with caution, since *β* also quantifies the overall goodness-of-fit of a model for an individual participant. If a particular model does not explain choices well for a participant, it will appear as though that participant is behaving more randomly with respect to the choice variables computed by the model (i.e., lower *β*). In the absence of other corroborating evidence, it is therefore not warranted to draw strong conclusions from the correlation between the *ϕ*_*cost*_ and *β* parameters.

### Principal Components Analysis

As previously described, we used a principal components analysis to quantify individual differences across common measures of each of three traits: need for structure/control, anxiety/negative emotionality, and obsessive-compulsion. In each of these principal component analyses, a clear one factor structure was indicated; we therefore extracted factor scores for the first principal component in each analysis as an estimate of the latent construct accounting for common variance across measures. For measures of the obsessive-compulsive trait, the first principal component accounted for 73% of the variance, and loadings were all greater than 0.8. For measures of the need for order and cleanliness trait, the first principal component accounted for 87% of the variance, with all loadings greater than 0.9. For measures of the anxiety/negative emotionality trait, the first principal component accounted for 72% of the variance, and loadings were all greater than 0.8.

### Relation Between Computational Model Parameters and Latent Self-Report Traits

In an important sense, our principal component analysis of questionnaire data and our computational modelling analysis of choice data serve the same purpose of interpretable dimensionality reduction. Both methods aim to extract psychologically meaningful latent factors from more complex raw data (individual scale responses on the one hand, and choice data on the other). With this in mind, we next sought to assess relations between preference for information and self-report constructs in the latent space defined by the computational model parameters and the extracted self-report factors. A Pearson correlation matrix summarising these associations (corrected for 20 comparisons using a False Discovery Rate correction; Benjamini & Hochberg, 1995) is presented in Table 4 (see Supplementary Figure 5 for scatterplots corresponding to each correlation).

**Table 4:** Pearson correlation matrix of computational model parameters and self-report factors and scales (N = 139).

|  | Need for<br>structure/control | Anxiety/negative<br>emotionality | Obsessive-compulsion | Intolerance of<br>uncertainty |
| --- | --- | --- | --- | --- |
| $\phi_{free}$ | .11 | -.01 | .05 | -.04 |
| $\phi_{cost}$ | .03 | .31 ** | .25 * | .15 |
| $k_{mean}$ | -.01 | -.02 | .05 | -.19<sup>†</sup> |
| $k_{var}$ | .12 | -.19<sup>†</sup> | -.05 | -.002 |
| $\log(\beta)$ | -.004 | -.16 | -.13 | -.15 |

\*\* $p < .01$ , corrected for multiple comparisons
\* $p < .05$ , corrected for multiple comparisons
<sup>†</sup> $p < .05$ , uncorrected

There was significant positive association between participants’ preference for costly information (quantified by the model parameter *ϕ*_*cost*_) and both anxiety/negative emotionality and obsessive-compulsion (see Figure 5). To contextualise the strength of these two associations, we note that meta-analyses estimate that an effect of *r =* 0.31 would fall within the upper quartile of effect sizes observed in individual-differences research (Gignac & Szodorai, 2016), and in the upper tertile of correlational effect sizes in behavioural science more broadly (Hemphill, 2003). The observed associations should therefore be considered relatively large.

**Figure 5.**
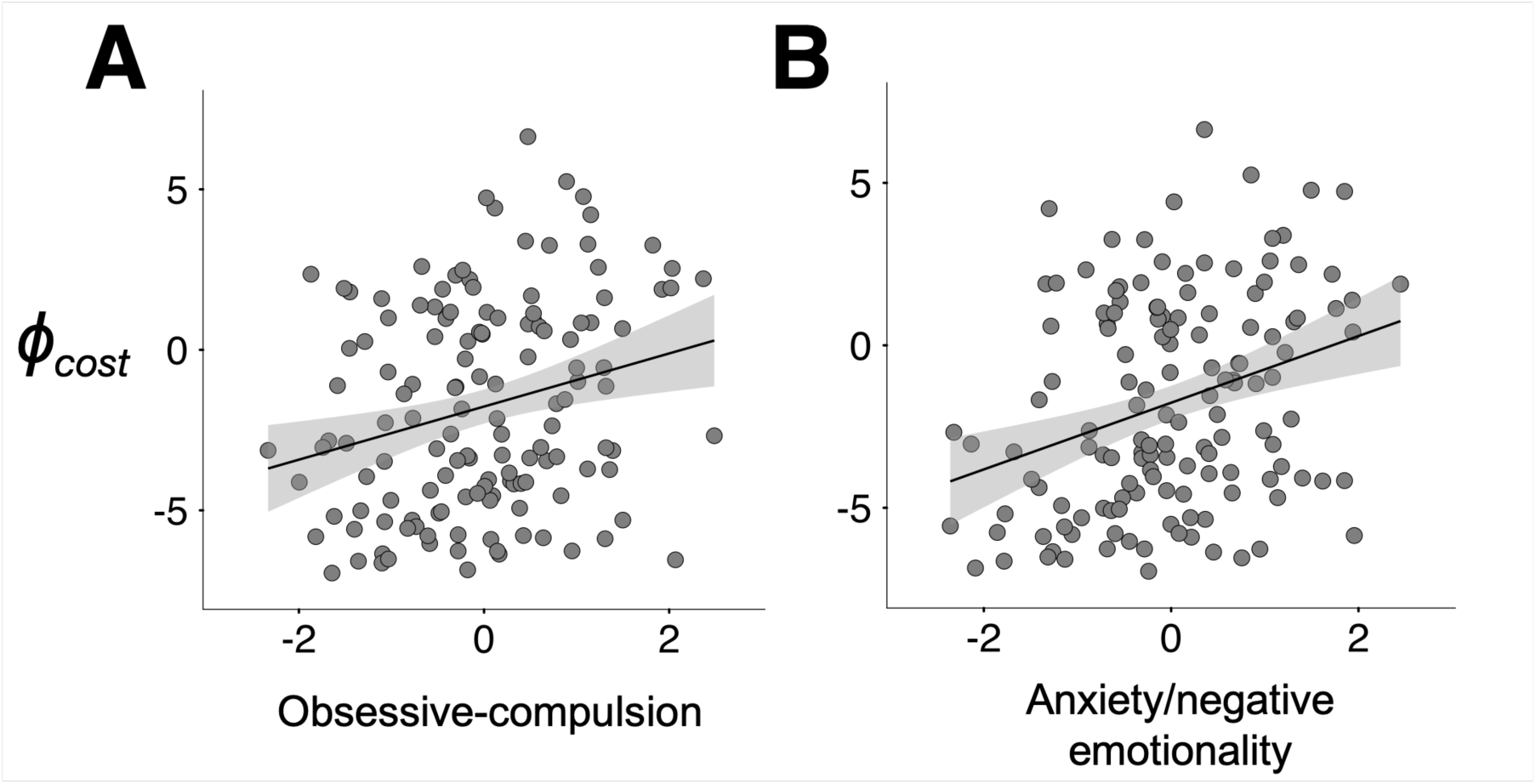
Scatterplot of parameter estimates and latent trait factor scores for individual participants. (A) Obsessive-compulsive traits were positively related to the *ϕ*_*cost*_ model parameter, which quantified willingness to pay for non-instrumental information. (B) The trait of anxiety/negative emotionality was also positively related to the *ϕ*_*cost*_ model parameter.

In addition, similar correlations were observed when preference for costly information was quantified using the second-best-fitting model (Model 11; see Supplementary Table 4), which was very similar to Model 13, but did not include the variance-dependent information preference parameter *k*_*var*_. Similar correlations were also observed when costly information preference was quantified using a model-agnostic metric (overall proportion of information-seeking choices in non-zero-cost conditions; see Supplementary Table 5). Taken together, these analyses further increase our confidence that the observed correlations were a meaningful pattern in participants’ choice data, and not an artefact of any particular measurement technique.

Partial correlation analyses suggested that the observed correlations were not likely to be due to covariance between the anxiety/negative emotionality and obsessive-compulsive factors. Each of the trait factors was significantly associated with *ϕ*_*cost*_ even when controlling for the other factor (correlation between *ϕ*_*cost*_ and obsessive-compulsive traits controlling for anxiety/negative emotionality: *r*(139) = 0.19, *p* < .05; correlation between *ϕ*_*cost*_ and anxiety/negative emotionality when controlling for obsessive-compulsive traits: *r*(139) = 0.27, *p* < .01). Indeed, a Fisher *r*-to*-z* transformation found no evidence that either partial correlation differed significantly from its non-partial counterpart (*p* > .6 for both). Our data are therefore consistent with the explanation that anxiety/negative emotionality and obsessive-compulsive traits accounted for unique variance in the distribution of the *ϕ*_*cost*_ parameter across participants.

Finally, two additional correlations—between *k*_*var*_ and anxiety/negative emotionality, and between *k*_*mean*_ and intolerance of uncertainty—were statistically significant, but did not survive correction for multiple comparisons.

Supplementary Table 3 presents a full correlation matrix between model parameters and individual self-report measures. These correlations demonstrate that the pattern of correlations observed between model parameters and latent self-report traits (Table 4) is a fair summary of the underlying relationships, giving us further confidence in our use of dimensionality reduction to quantify latent self-report traits.

## Discussion

This study examined associations between subclinical obsessive-compulsive and anxious traits and preference for acquiring non-instrumental information. We collected rich self-report and behavioural data and used dimensionality reduction techniques to quantify psychologically interpretable latent variables within each dataset. For self-report data, we performed dimension reduction via a principal component analysis, and extracted measures of three latent traits: need for structure/control, anxiety/negative emotionality, and obsessive-compulsion. For choice data, we first developed a number of computational models that could account for individual differences in information-seeking behaviour in the gain domain, the loss domain and a mixture of both. Then, using a hierarchical Bayesian modelling approach, we identified a best-fitting computational cognitive model that included 5 separate parameters per participant, each corresponding to a distinct variable in the decision process. By estimating each of these parameters for every participant, we were able to quantify the degree to which participants differed in the different components of the choice process.

We found significant positive correlations between the computational model parameter that quantified participants’ valuation of costly information (*ϕ*_*cost*_) and both anxiety/negative emotionality and obsessive-compulsion. These correlations were relatively large (Gignac & Szodorai, 2016; Hemphill, 2003) and statistically significant, even when controlling for one another, indicating that each trait explained unique variance in participants’ willingness to pay for non-instrumental information. Interestingly, there was no evidence for a significant association between any latent trait and a model parameter quantifying participants’ preference for costless information (*ϕ*_*free*_), and no evidence that any trait was related to modulations of information preference across different lottery payout domains. We therefore conclude that the primary behavioural phenotype associated with anxious and obsessive-compulsive traits in the present study was a willingness to incur a cost in order to reduce uncertainty about a future event.

These results suggest a link between aversion to uncertainty—manifesting behaviourally as increased willingness to pay for non-instrumental information—and both anxious and obsessive-compulsive traits. In recent years, theorists have increasingly advocated dimensional perspectives for psychiatric disorders including OCD and generalised anxiety. On this account, subclinical traits such as those measured in the present study lie on a continuum with psychiatric symptoms experienced by individuals who meet full diagnostic criteria for a disorder (Haslam et al., 2012; Robbins et al., 2012; Widiger et al., 2019). Thus, our findings offer support for psychiatric theories that implicate affective responses to uncertainty about the future as a cognitive risk factor for obsessive-compulsive and anxiety disorders (Carleton et al., 2012; McEvoy & Erceg-Hurn, 2016; Miceli & Castelfranchi, 2005; Shihata et al., 2016). In the case of OCD, cognitive-behavioural theories propose that compulsions develop in OCD because they reduce subjective distress regarding aversive thoughts, and are thereby reinforced (Carr, 1974; Rachman, 2002; Salkovskis, 1985). In particular, checking behaviours can be conceptualised as having an uncertainty-reduction goal (for example, repeatedly checking the knobs on a gas stove to reduce one’s uncertainty about the possibility that the stove has been left on). According to this explanation, individuals with high trait-level aversion to uncertainty would be more vulnerable to developing a checking compulsion because the relief from uncertainty engendered by checking would be more reinforcing, and therefore more likely to be repeated (Lind & Boschen, 2009). Compulsive checking might therefore be seen as an information-seeking behaviour that has been negatively reinforced to the extent of becoming habitual (see, e.g., Gillan et al., 2014). Our results also provide support for theories linking aversion to uncertainty with cognitive symptoms of anxiety disorders (Freeston et al., 1994; Koerner & Dugas, 2006; Miceli & Castelfranchi, 2005). In particular, our results are consistent with a process model of anxiety in which uncertainty about the future is aversive because it is perceived as a lack of epistemic control (Miceli & Castelfranchi, 2005). On this account, individuals with anxiety are characterised by a desire for absolute predictive certainty about the future, such that uncertainty of any degree signals an intolerable lack of control. On this assumption it is rational to seek out and pay for non-instrumental information, which delivers certainty about a future outcome even if it does not allow one to alter that outcome. Further research in psychiatric patient samples is necessary to test this hypothesis.

Interestingly, participants’ choices revealed a significant anticorrelation between preference for costless information and preference for costly information. This pattern of behaviour has not been previously documented, but re-analysis of published datasets from the non-instrumental information-seeking task revealed a similar pattern of results (Bennett et al., 2016; Brydevall et al., 2018). This suggests that willingness to acquire costless information and willingness to pay for costly information might be dissociable behavioural phenotypes in humans. Of course, since we observed a negative correlation between these two measures, it would not be accurate to describe them as strictly independent of one another. Nevertheless, the observed negative association is inconsistent with the possibility that the two measures are manifestations of the same latent construct (in which case we would instead expect to observe a *positive* association).

A dissociation between preference for free information and preference for costly information is in contrast to typical assumptions and may help to explain some apparent inconsistencies in previous research. For instance, the standard version of the information sampling task as implemented in the CANTAB neuropsychological battery (Clark et al., 2006) includes both a costless information measure (‘fixed win’ condition) and a costly information measure (‘decreasing win’ condition). Both measures are treated as measures of the same latent construct; however, observed effects are frequently specific to either the fixed win condition or the decreasing win condition (Clark, Robbins, Ersche, & Sahakian, 2006; Crockett, Clark, Smillie, & Robbins, 2012; Mole et al., 2015; Valls-Serrano, Caracuel, & Verdejo-Garcia, 2016; but see also Bennett et al., 2017; Bennett, Yücel, & Murawski, 2018). Similarly, research using the beads task has shown that the classic ‘jumping-to-conclusions’ bias ((whereby participants with psychosis request less information prior to making a decision than healthy controls; Huq et al., 1988) is reversed when information is costly rather than free (Baker et al., 2019). A dissociation between preferences for costless and costly information may help to account for these apparent discrepancies. In this respect, it is noteworthy that participants’ self-reported curiosity about future lottery outcomes has been found to be proportional to willingness to pay a wait-time cost to observe the outcome of this lottery (Jach & Smillie, in press; van Lieshout et al., 2018). This suggests the possibility that preference for costly information is more strongly related to subjective feelings of curiosity than preference for costless information; this hypothesis should be investigated in future research.

Our finding that anxious and obsessive-compulsive traits were related to preference for costly information, but not costless information, suggests a more nuanced account of the link between aversion to uncertainty and psychopathology. Our results suggest that preference for costless information is relatively common, but that a willingness to acquire information when it comes at a considerable cost is much rarer, and linked with anxious and obsessive-compulsive traits. This variability in the change of preference with changes in cost is captured by the microeconomic concept of price elasticity. Our results are therefore consistent with a specific association between obsessive-compulsive and anxious traits and an *inelastic* preference for information. In OCD or generalized anxiety, this could manifest in symptoms such as compulsive checking and reassurance-seeking, which involve incurring significant social, personal, or opportunity costs in an attempt to abolish uncertainty (Kobori & Salkovskis, 2013; Salkovskis, 1985). In our computational model, variable price elasticity between participants was captured by an information preference parameter that varied categorically between costless information trials and costly information trials. We used this modelling approach, which outperformed a separate approach assuming variability in linear price elasticity across participants, because we acquired data for only three datapoints on the non-instrumental information demand curve (0, 1, and 3 cents); as such, we did not have sufficient data to infer the parametric form of the price elasticity function.

At a group level, we also found that participants preferred information about lotteries in the loss domain less than information about lotteries in either the gain domain or the mixed domain, consistent with a previous study (Charpentier et al., 2018). Our computational modelling results therefore provide further evidence that preference for non-instrumental information about a lottery varied in proportion to both mean and variance of lottery outcomes, in line with a recent study by Kobayashi et al. (2019). Participants’ preference for information was still positive for lotteries in the loss domain, meaning that the present study did not find any evidence for information *avoidance*; however, this phenomenon has been documented in more naturalistic decision problems, especially for possible outcomes that are extremely negatively valenced (e.g., medical tests) (Golman et al., 2017; Sweeny et al., 2010). These findings are beyond the scope of the present paper, and further research is required to build a complete computational model of information avoidance. One possible explanation is suggested by decision affect theory, which proposes that decision-makers act on preferences over their expected emotional response to future events (Mellers, 2000; Mellers et al., 1999). This framework can predict information avoidance for very negative outcomes in the loss domain as the result of participants’ anticipation of a negative emotional response to early information (see also Sharot & Sunstein, 2020). In line with this interpretation, Kobayashi et al. (2019) recently showed that when participants could choose to obtain non-instrumental information about one of two lotteries, they showed a strong preference for acquiring information about the lottery with the highest expected value (even after accounting for the expected information gain). <u>The authors interpreted these findings in terms of the affective consequences of advance information about positive and negative outcomes</u> (i.e., savouring wins in the gain domain; avoiding dread of losses in the loss domain). Interestingly, however, the expected-value-related preference for information in this study was not related to anxious or obsessive-compulsive traits. This stands in contrast to the association between these traits and preference for costly information as measured in the present study, but is consistent with our finding that the *k*_*mean*_ parameter was uncorrelated with either anxiety or obsessive-compulsion.

In this respect, one limitation of the present study is the relatively restricted range of payoff conditions tested (i.e., three win/loss outcome pairs: +20c/0c, +20c/-20c, 0c/-20c), which may limit the interpretability of the mean- and variance-dependent information preference parameters *k*_*mean*_ and *k*_*var*_. Specifically, a positive *k*_*mean*_ parameter is consistent with two possible interpretations: on the one hand, information preference might be proportional to the mean value of the lottery; however, it is also possible that information preference might increase in strength only for lotteries that do not include a loss outcome. We favour the former interpretation, since the positive overall value of the *k*_*var*_ parameter indicates that preference for information was strongest overall in the high-variance (+20c/-20c) lottery condition, contrary to what would be predicted if information preference was determined by the presence or absence of a loss outcome. This interpretation is also consistent with the results of Kobayashi et al. (2019), who showed that preference for information about a gamble was proportional to the expected value of the gamble across both the gain domain and the loss domain. Future research could further investigate this question by using a more varied set of lottery outcome combinations, which would increase the interpretability of model parameters.

We did not find strong evidence for an association between any aspect of participants’ choice behaviour and scores on the Intolerance of Uncertainty scale. This result is contrary to what might be expected given the theoretical link between information-seeking and aversion to uncertainty, and suggests two possible interpretations. First, behavioural and self-report measures might capture distinct variance in the underlying construct of aversion to uncertainty, as has been reported for other individual-difference constructs such as impulsivity (Cyders & Coskunpinar, 2012). Second, although we designed this study with sufficient power to detect a typical individual-differences effect size (see Method) (Fraley & Marks, 2007), we cannot rule out the possibility of weaker associations between self-report and behavioural assays of aversion to uncertainty. We note that results revealed a small negative correlation between self-reported intolerance of uncertainty and the *k*_*cost*_ model parameter, but that this effect did not survive correction for multiple comparisons. Further research with better power to detect small associations would be required to provide conclusive evidence regarding this question.

Looking forward, the field awaits a computational psychiatric model (Bennett et al., 2019; Huys et al., 2016) of the role played by aversion to uncertainty in disorders like generalised anxiety and OCD. Many details of this theoretical framework are yet to be identified, key among them the cognitive mechanism by which particular subjects of uncertainty come to take on especial salience in these disorders. This is critical, because at any given moment there is an infinite number of future events about which an individual with anxiety or OCD is uncertain, and uncertainty about the vast majority of these is not in the least aversive. The results of present study suggest that, conditional on attention having been directed to a future lottery, those who self-report more anxious and obsessive-compulsive traits find their uncertainty to be more aversive, and are therefore more likely to incur a monetary cost to acquire information. However, this result is mute on the question of how future outcomes acquire salience in natural settings. One possibility that might be tested in future research is that, given the well-documented attention biases to threat in anxiety disorders (MacLeod & Mathews, 1988), aversion to uncertainty might interact with attention capture by potential future threat in anxiety and OCD.

In summary, we find that two distinct subclinical traits—obsessive-compulsion and anxiety/negative emotionality—are independently related to valuation of non-instrumental information in a general population sample. This finding is consistent with theories proposing aversion to uncertainty as a dimensional transdiagnostic trait underlying psychopathology. Our results specifically identify willingness to pay for non-instrumental information as a candidate behavioural phenotype for anxiety and OCD, and future research testing this preference in psychiatric patient samples will be of great interest. Simultaneously, our results extend previous computational cognitive studies of information-seeking by showing that human preferences for non-instrumental information are not invariant across all sources of uncertainty. Instead, preference for non-instrumental information appears to be modulated by several distinct properties of the object of uncertainty, including (but likely not limited to) the mean and variance of the utilities of possible future events.

