## Supplementary Materials for "Anxious and obsessive-compulsive traits are independently associated with valuation of non-instrumental information"

**Supplementary Information**

*Supplementary Table 1: Spearman correlation matrix of overall information preference (quantified as proportion of information-seeking choices) across different payout domains.*

| | Gain domain | Loss domain | Mixed domain |
|---|---|---|---|
| Gain domain | 1 | - | - |
| Loss domain | .74 *** | 1 | - |
| Mixed domain | .87 *** | .78 *** | 1 |

*** $p < .001$

*Supplementary Table 2: Pearson correlation matrix of computational model parameters and individual scale totals*

| | Obsessive-compulsive traits | | | Need for structure/ control | | Anxiety/negative emotionality | | | |
|---|---|---|---|---|---|---|---|---|---|
| | Obsessive-compulsive inventory | Rigid perfectionism | Need for order and cleanliness | BFI-2 organisation | BFAS orderliness | BFI-2 anxiety | BFI-2 emotional volatility | BFAS withdrawal | BFAS volatility |
| Obsessive-compulsive inventory | 1 | - | - | - | - | - | - | - | - |
| Rigid perfectionism | .66 | 1 | - | - | - | - | - | - | - |
| Need for order and cleanliness | .47 | .65 | 1 | - | - | - | - | - | - |
| BFI-2 organisation | .15 | .39 | .74 | 1 | - | - | - | - | - |
| BFAS orderliness | .25 | .49 | .79 | .75 | 1 | - | - | - | - |
| BFI-2 anxiety | .28 | .23 | .13 | .01 | .18 | 1 | - | - | - |
| BFI-2 emotional volatility | .26 | .19 | .00 | -.24 | -.05 | .55 | 1 | - | - |
| BFAS withdrawal | .25 | .19 | .06 | -.09 | .09 | .69 | .59 | 1 | - |
| BFAS volatility | .31 | .23 | .06 | -.16 | .02 | .57 | .77 | .60 | 1 |

Shaded blue areas denote within-factor correlations. For correlations in this matrix, the threshold for statistical significance at $\alpha = .05$ is $r = \pm .17$.

*Supplementary Table 3: Pearson correlation matrix of computational model parameters and individual scale totals (N = 139).*

| | Obsessive-compulsive traits | | | Need for structure/control | | Anxiety/negative emotionality | | | |
|---|---|---|---|---|---|---|---|---|---|
| | Obsessive-compulsive inventory | Rigid perfectionism | Need for order and cleanliness | BFI-2 organisation | BFAS orderliness | BFI-2 anxiety | BFI-2 emotional volatility | BFAS withdrawal | BFAS volatility |
| $\phi_{free}$ | .05 | -.01 | .08 | .13 | .07 | .07 | -.01 | -.07 | -.02 |
| $\phi_{cost}$ | .24 ** | .25 ** | .15 | -.01 | .06 | .23 ** | .27 ** | .25 ** | .32 *** |
| $k_{mean}$ | .03 | .06 | .04 | .03 | -.04 | -.09 | -.01 | .03 | -.01 |
| $k_{var}$ | -.09 | -.09 | .06 | .11 | .10 | -.12 | -.18 * | -.18 * | -.18 * |
| $\log(\beta)$ | -.10 | -.13 | -.10 | .0003 | -.01 | -.10 | -.15 | -.15 | -.15 |

*** $p < .001$; ** $p < .01$; * $p < .05$

*Supplementary Table 4: Pearson correlation matrix of computational model parameters and self-report factors and scales for Model 11 (N = 139).*

| | Need for structure/control | Anxiety/negative emotionality | Obsessive-compulsion | Intolerance of uncertainty |
|---|---|---|---|---|
| $\phi_{free}$ | .11 | -.02 | .04 | -.03 |
| $\phi_{cost}$ | .03 | .31 ** | .25 * | .14 |
| $k_{mean}$ | -.001 | -.03 | .06 | $-.18^{\dagger}$ |
| $\log(\beta)$ | -.02 | -.14 | -.12 | -.16 |

** $p < .01$, corrected for multiple comparisons

* $p < .05$, corrected for multiple comparisons

$^{\dagger}p < .05$, uncorrected

*Supplementary Table 5: Pearson correlation matrix of information choice proportions and self-report factors and scales (N = 139).*

| | Need for structure/control | Anxiety/negative emotionality | Obsessive-compulsion | Intolerance of uncertainty |
|---|---|---|---|---|
| Preference for free info | .11 | .004 | .04 | -.01 |
| Preference for costly info | .04 | .24 ** | .28 *** | .12 |

*** $p < .001$
** $p < .01$

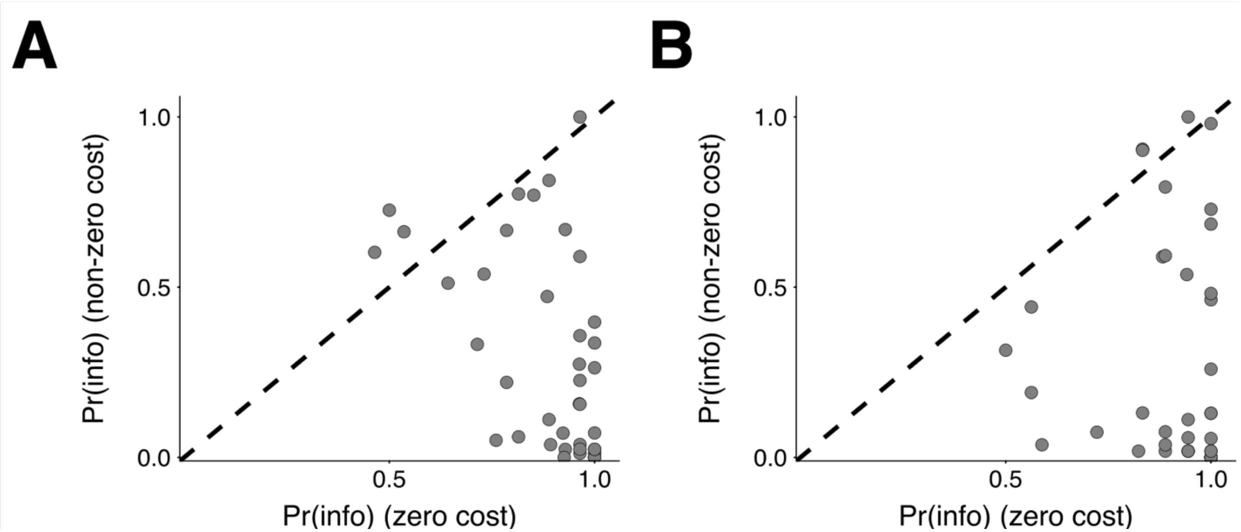


*Supplementary Figure 1*: Re-analysis of previously collected data using the non-instrumental information-seeking task revealed a pattern of results in line with the present study. (A) In data ($n$ = 40) collected by Bennett et al. (2016; *PLoS Computational Biology*) there was a significant negative correlation between preference for information in the zero-cost condition and mean preference for information in non-zero cost conditions (Spearman $\rho$ = -.51, $p < .01$). (B) In data ($n$ = 40) collected by Brydevall et al. (2018, *Scientific Reports*), there was a non-significant trend towards a negative correlation between these two quantities (Spearman $\rho$ = -.30, $p$ = .06).

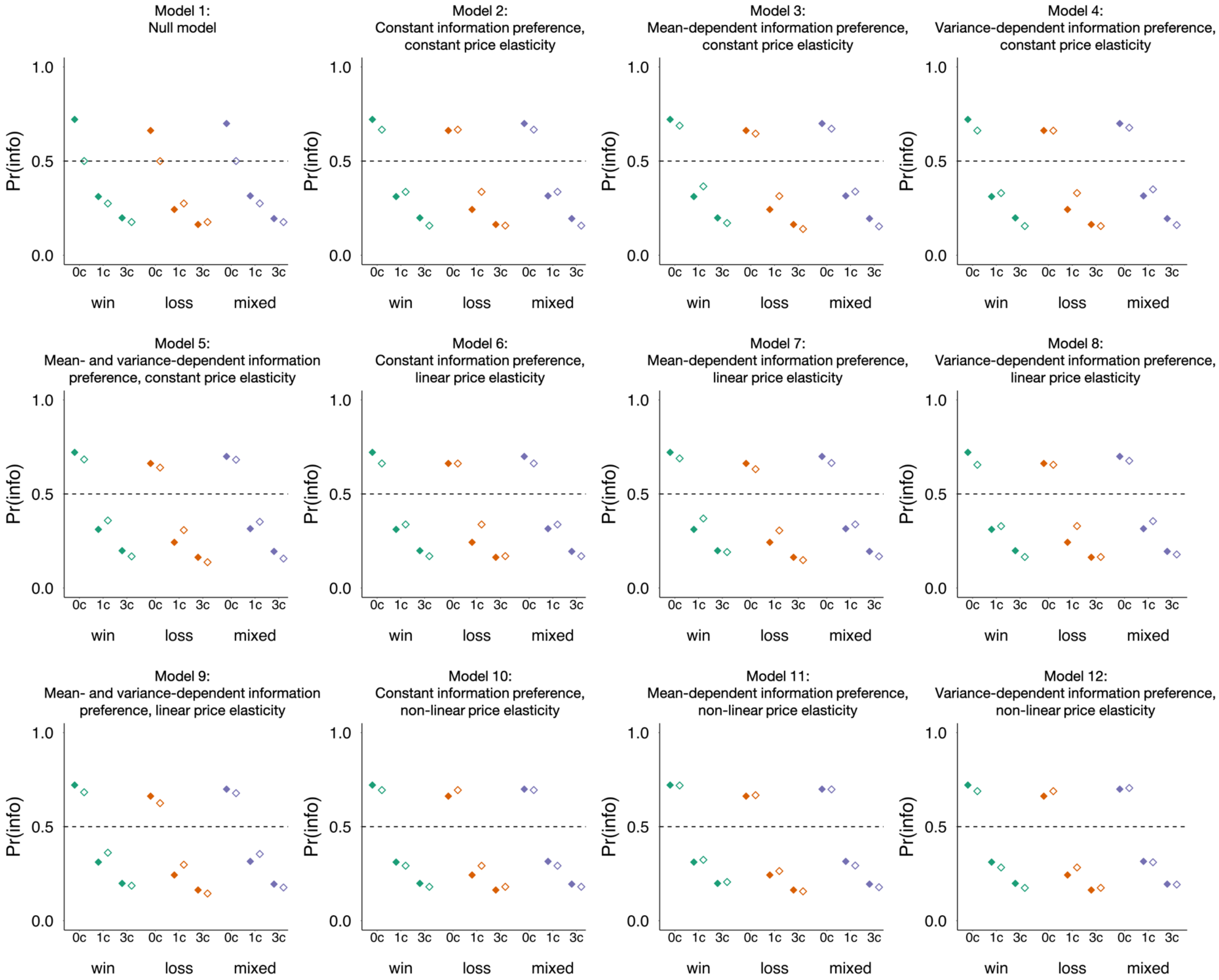


*Supplementary Figure 2*: Posterior predictive checks for the 12 rejected models. Data are observed (filled markers) and predicted (unfilled markers) mean informative-stimulus choice proportions across payout domains and cost conditions.

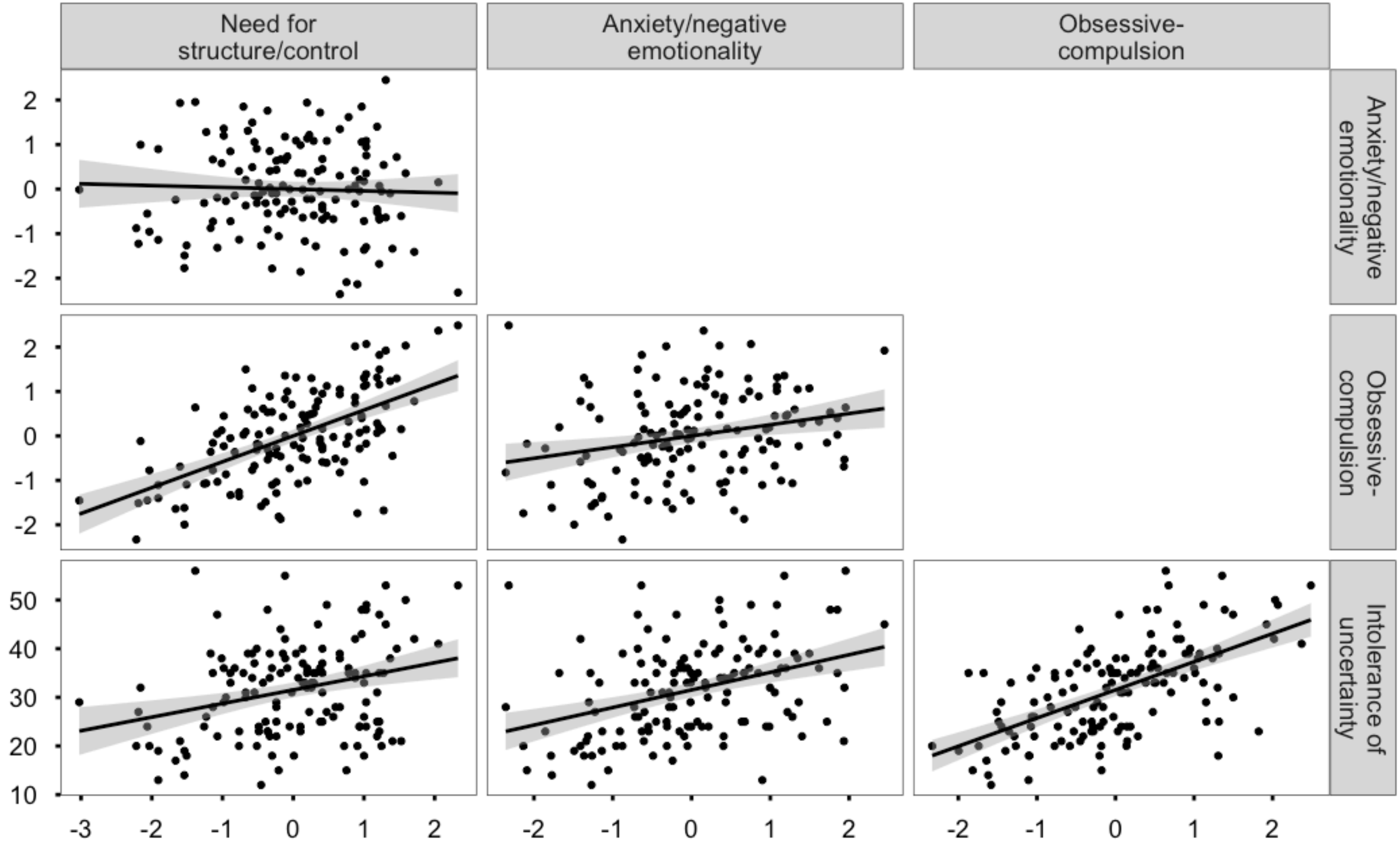


*Supplementary Figure 3*: Scatterplots of extracted factors and scales in self-report battery (corresponding to correlations reported in Table 1 of the manuscript). Each dot represents a factor or scale score for one participant. Shaded areas represent the 95% confidence interval of the respective lines of best fit.

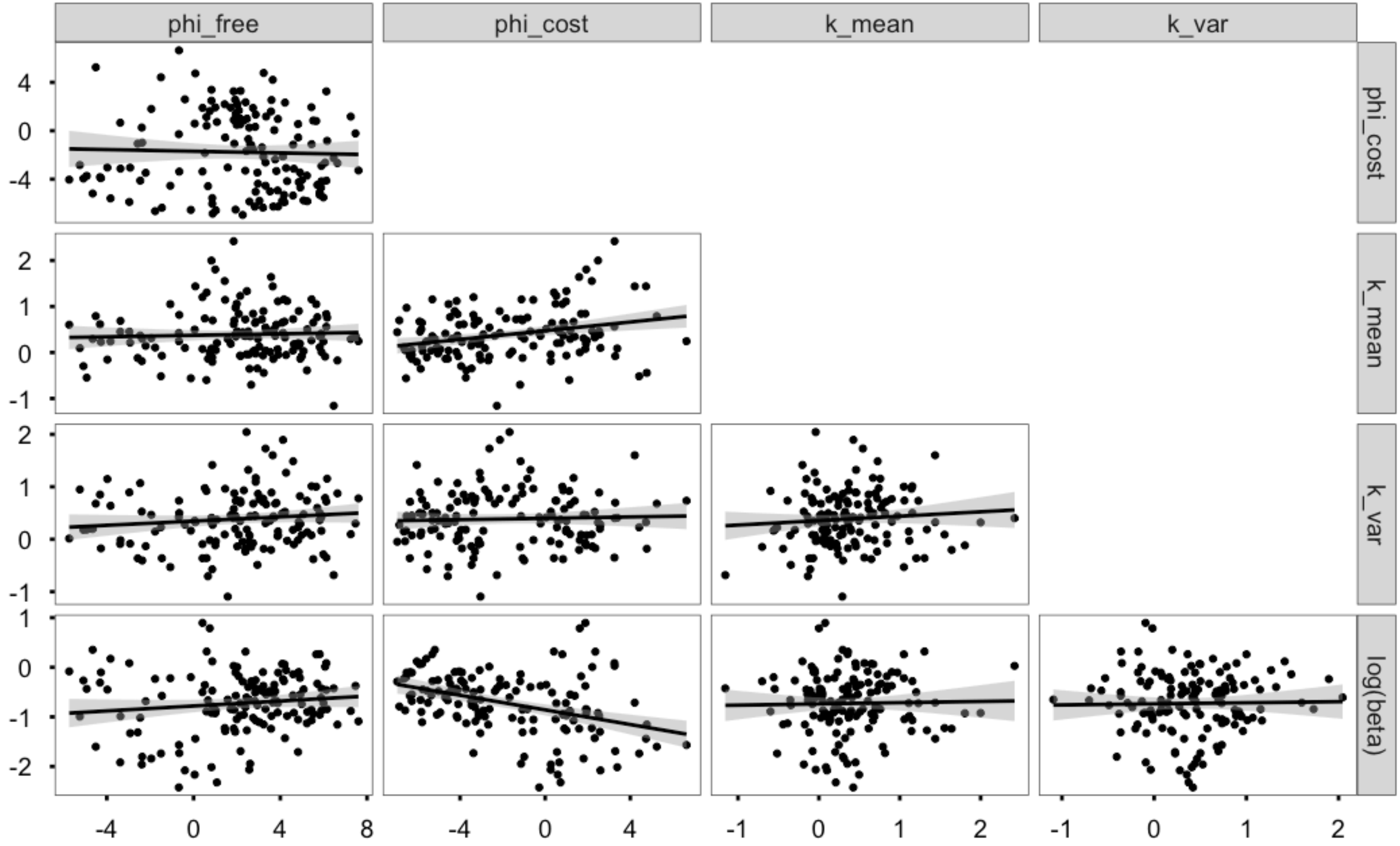


*Supplementary Figure 4*: Scatterplots of estimated model parameters across participants (corresponding to correlations reported in Table 3 of the manuscript). Shaded areas represent the 95% confidence interval of the respective lines of best fit.

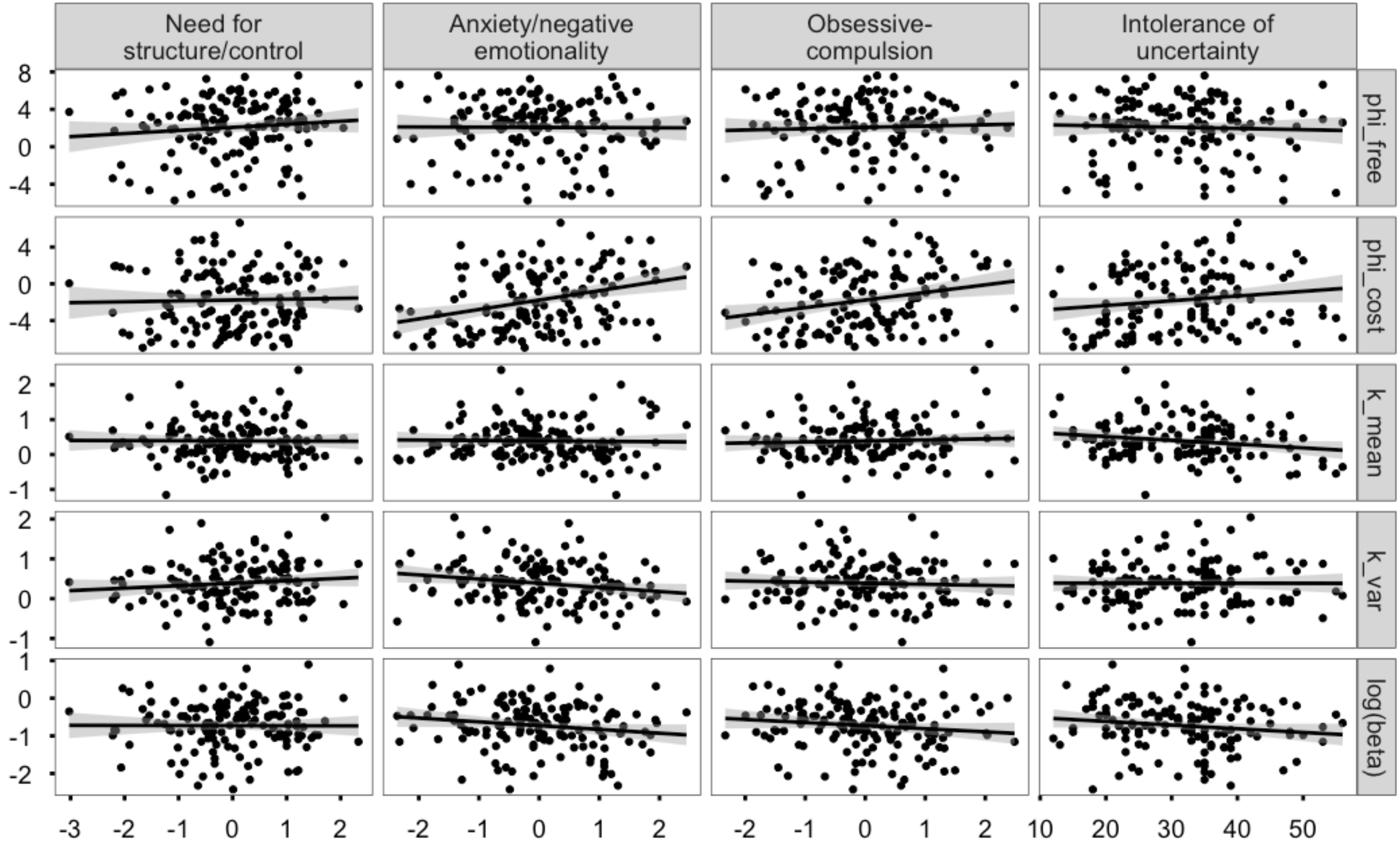


*Supplementary Figure 5*: Scatterplots for self-report factors (columns) and model parameter estimates (rows). Subplots in this figure correspond to the correlation matrix reported in Table 4 of the manuscript. Each dot represents a factor or scale score for one participant. Shaded areas represent the 95% confidence interval of the lines of best fit.
